# Towards a better understanding of the low recall of insertion variants with short-read based variant callers

**DOI:** 10.1101/2020.06.09.142232

**Authors:** Wesley Delage, Julien Thevenon, Claire Lemaitre

## Abstract

Since 2009, numerous tools have been developed to detect structural variants (SVs) using short read technologies. Insertions >50 bp are one of the hardest type to discover and are drastically underrepresented in gold standard variant callsets. The advent of long read technologies has completely changed the situation. In 2019, two independent cross technologies studies have published the most complete variant callsets with sequence resolved insertions in human individuals. Among the reported insertions, only 17 to 37% could be discovered with short-read based tools. In this work, we performed an in-depth analysis of these unprecedented insertion callsets in order to investigate the causes of such failures. We have first established a precise classification of insertion variants according to four layers of characterization: the nature and size of the inserted sequence, the genomic context of the insertion site and the breakpoint junction complexity. Because these levels are intertwined, we then used simulations to characterize the impact of each complexity factor on the recall of several SV callers. Simulations showed that the most impacting factor was the insertion type rather than the genomic context, with various difficulties being handled differently among the tested SV callers, and they highlighted the lack of sequence resolution for most insertion calls. Our results explain the low recall by pointing out several difficulty factors among the observed insertion features and provide avenues for improving SV caller algorithms and their combinations.

## Introduction

The widespread use of short read massively parallel sequencing has allowed the fine characterization of the human genome variability on single nucleotide variants and small insertions/deletions (<50 bp) [1, 2]. Structural variants (SVs) are larger variants. They are defined as a fragment of DNA of more than 50 bp that differs between an individual and the reference genome [3]. There is a great variety of SVs, with various proposed stratifications. A common categorisation differentiates a deletion (DEL) for a loss of a fragment, an insertion (INS) for a gain of a fragment, an inversion for a reversion of a fragment (INV) and a translocation (TRANS) for moving a fragment to another position in the genome. SVs are drivers of the genome evolution along generations, and some of them can have a significant functional impacts on the organism and be responsible for rare Mendelian disorders [4].

The classical approach to discover SVs from Whole Genome Sequencing (WGS) with short reads relies on a first step consisting in mapping the reads to a reference genome. Then SV callers look for atypical mapping signals, such as discordant read pairs, clipped reads or abnormal read depth, to identify putative SV breakpoints along the reference genome [5, 6]. More than 70 SV callers have been developed up to date and several benchmarks have revealed great variability between results obtained by different methods, demonstrating that SV detection using short read sequencing remains challenging [7, 8]. The challenge is to resolve two issues: a technical and a methodological one. The technical issue concerns the sequencing technology: insert size, read size and sequencing coverage have been shown to impact SV discovery [9]. The second issue concerns SV caller algorithms and their ability to decipher and translate the biological signal from the alignments. Thus, SVs located in repeated regions or containing repeats larger than the read size are difficult to detect [9].

In particular, insertions are one of the most difficult SV types to call [8, 7]. Because the inserted sequence is absent from the reference genome, or at least at the given locus of insertion, calling such variants and resolving the exact inserted sequence requires finely tuned approaches such as *de novo* or local assembly [10, 11]. This increased difficulty is well exemplified by the dramatic under-representation of such SV type in usual reference databases or standard variant callsets. For instance, dbVar at present references only 28 % of insertions or duplications among the SVs larger than 50 bp. On the opposite, deletions represent more than 70 % of the database, although both types are expected to be roughly equally abundant in human populations[12]. Moreover, only 1.5% of the reported insertions are sequence-resolved, that is with an inserted sequence fully characterized.

One explanation is that the size of the reads is small compared to the target event size and the detection is mainly based on alignments which may produce artefacts[13]. Another source of difficulty for insertion detection is the presence of repeated patterns at the precise rearrangement breakpoints. Several molecular mechanisms involved in rearrangement genesis are known to produce such repeated sequences, referred as junctional homology [14, 15, 16]. Junctional homology is defined as a DNA sequence that has two identical or nearly identical copies at the junctions of the two genomic segments involved in the rearrangement, when the sequence is short (< 70 bp) this is often called a micro-homology [16]. The repair of DNA double strand breaks by diverse mechanisms, such as Non-Allelic Homologous Recombination (NAHR), Non-Homologous End Joining (NHEJ) or Microhomology-Mediated Break-Induced Replication (MMBIR), generate such homologies whose size depend precisely on the type of the involved mechanism. These homologies can have an impact on insertion calling performance, since the concerned region at the inserted site is no longer specific to the reference allele and it is no longer possible to identify the exact location of the insertion site. However, little is known at present about the prevalence of these homologies and their sizes for human insertion variants due to their poor referencing in databases.

More recently, novel long reads sequencing technologies have overcome these limitations and allowed the generation of more accurate datasets, finally referencing sequence-resolved insertion variants in the human genome[17, 8]. Thanks to several international efforts, some gold standard callsets have been produced in 2019, referencing tens of thousands of insertions in several human individuals [18, 19]. Among the reported insertions by Chaisson et al, a great majority (83 %) could not be discovered by any of the tested short-read based SV callers. This result of recall below 17 % is drastically different from the announced performances of insertion callers when evaluated on simulated datasets [20]. Indeed, Chaisson et al showed that 59 % of insertion variants were found in a tandem repeat context, suggesting that most of the real insertion variants in human individuals are probably occurring in complex regions and involving complex sequences. So far, such complexity factors were rarely included nor analysed in method benchmarks and to do so, actual insertion variants require to be better characterized.

Numerous countries are developing genomic medicine programs, based on short-read sequencing. Although third generation sequencing offers an unprecedented technique for exploring the complexity of individual structural variants, most of the genomic sequencing facilities will still use short-read based sequencing in coming years for its reduced cost. Hence, there is a critical need to measure and control the caveats of standard procedures for detecting SVs with short-read sequencing data.

In this work, we performed an in-depth analysis of these unprecedented insertion callsets, in order to investigate the causes of short read based caller failures. We have first established a precise classification of insertion variants according to four different layers of characterization: the nature and size of the inserted sequence, the genomic context of the insertion site and the breakpoint junction complexity. Because these levels are intertwined, we then used simulations to characterize the impact of each complexity factor on the recall of several SV callers.

## Results

### In-depth analysis of an exhaustive insertion variant callset

In this work, we first aimed at precisely characterizing an exhaustive set of insertion variants present in a given human individual. We based our study on a recently published SV callset published by Chaisson and colleagues in 2019 [18]. Using extensive sequencing datasets, combining different sequencing technologies and methodological approaches (short, linked and long reads, mapping-based and assembly-based SV calling), three human trios were thoroughly analysed to establish exhaustive and gold standard SV callsets (Supplementary Table 1). We first focused our study on the individual NA19240, son of the so-called Yoruban (YRI) Nigerian trio, whose SV callset contained 15,693 insertions greater than 50 bp.

#### Nature and size of the inserted sequences

Insertion variants can be classified in different sub-types according to the nature of the inserted sequence. Three insertion categories were distinguished in the original publication, namely tandem repeats, mobile element insertions and complex ones for all the other types. We proposed to refine this classification in five insertion sub-types, illustrated in Figure 1. A classical subdivision consisted in distinguishing *novel sequence* insertions from insertions of exiting sequences, namely duplicative insertions. Several sub-types of duplicative insertions were then defined according to the location or amount of the inserted sequence copies in the reference genome. Among duplicative insertions, we proposed to stratify (i) *tandem duplications*, with at least one copy of the inserted sequence being adjacent to the insertion site, (ii) *dispersed duplications*, with copies that can be located anywhere else in the genome. Among tandem duplications, *tandem repeats* are characterized by multiple tandem repetitions of a seed motif within the inserted sequence. *Mobile element* insertions are a very specific sub-type whose sequences are known and referenced in families. They are notably characterized by very high copy numbers in the genome (typically greater than 500). Other dispersed duplication types were then required to have a copy number lower than 50, in order not to be confounded with potential mobile element insertions.

**Figure 1:**
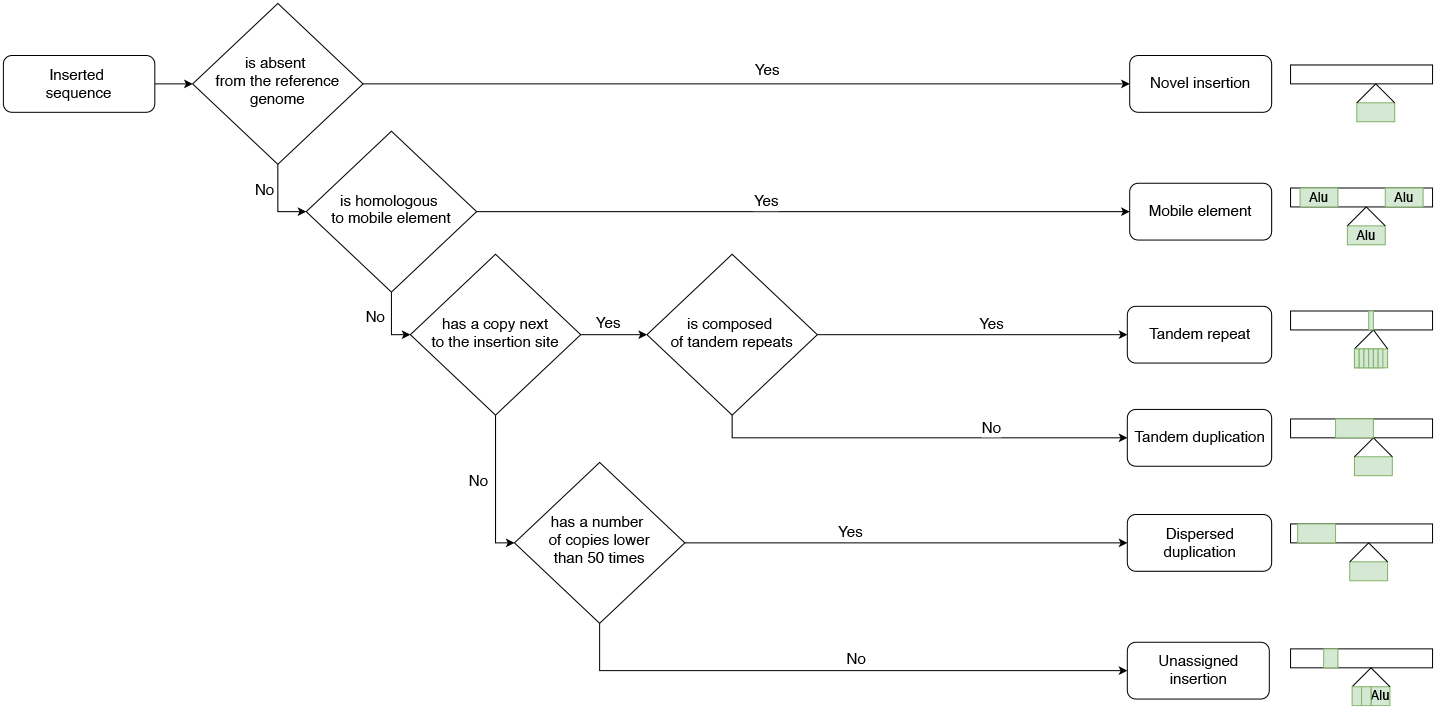
Decision tree used to classify insertion variants. Five insertion sub-types are defined according to the nature of the inserted sequence: novel sequence, tandem repeat and mobile element insertions and tandem and dispersed duplications. Unassigned insertions contain insertions which not meet requirements to be assigned to at least one sub-type.

In order to classify the insertion callset, all inserted sequences were aligned against the human reference genome, a mobile element database and were scanned for tandem repeats (see Material and Methods). We used a minimal sequence coverage threshold to annotate each insertion to an insertion sub-type according to the decision tree described in Figure 1. Insertions that did not meet any requirement to be annotated as one of the previous sub-types were qualified as *unassigned* insertions.

We set the threshold to 80% for our analysis to ensure a compromise between specificity and quantity of annotated insertions in all sub-types. With such threshold, 88% of insertions could be assigned to a given type. Among the 15,693 insertions, 8,735 (56%) were annotated as tandem repeats, 2,473 (16%) as mobile elements, 1,000 (6%) as tandem duplications, 869 (6%) as novel sequences and 773 (5%) as dispersed duplications (Figure 2 B and Supplementary Table 2 for results obtained with other coverage thresholds). 46% of tandem repeats had a repeat seed smaller than 10 bp and 93% smaller than 50 bp. Compared to the classification of Chaisson et al, the proportions of tandem repeats (57% vs 56%) and mobile elements (23% vs 16%) were close. The difference in mobile element proportions mainly represented insertions that were unassigned in our annotation. The 1,843 (12%) unassigned insertions at 80% threshold showed partial annotations of mobile element (57%), tandem repeats (22%), tandem duplications (15%) or dispersed duplications (5%).

**Figure 2:**
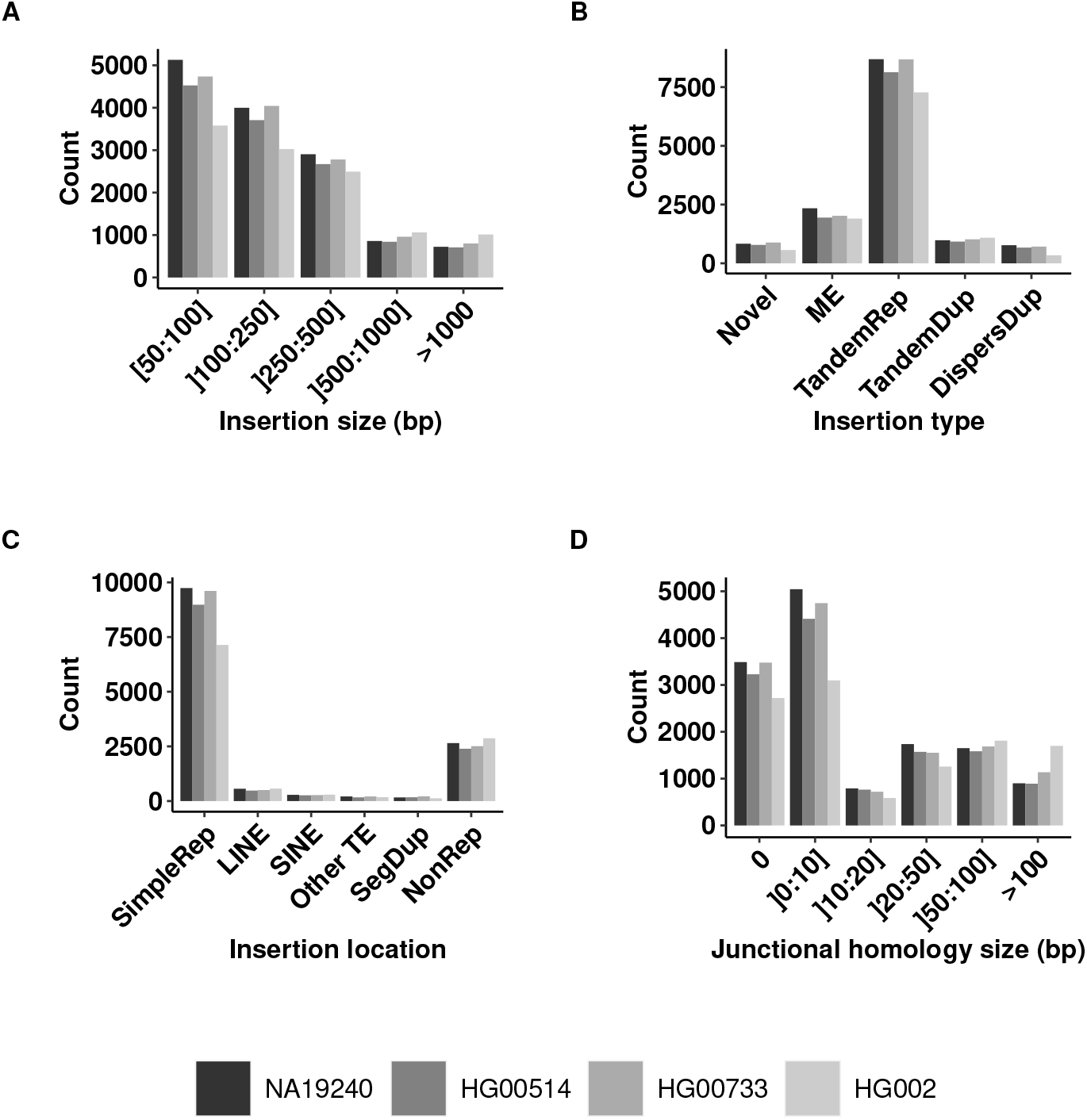
Distributions of insertion variant features across several callsets. Distributions of (A) insertion size, (B) insertion type, (C) repeated context of insertion and (D) homology size at the breakpoint for NA19240, HG00514, HG00733 and HG002 insertion variant callsets. Abbreviations: SimpleRep for simple repeat, ME for mobile element, TandemRep for tandem repeat, TandemDup for tandem duplication, DispersDup for dispersed duplication.

Concerning the size of the insertions, 67% of the insertions were smaller than 250 bp and only 8 % had a size greater than 1 Kb (Figure 2 A). Interestingly, the size distributions differed between insertion types (Figure 3 A). Mobile elements showed the most contrasting size distribution with a strong over-representation of the 250-500 bp size class (61 %). This can be explained by the most frequent and active mobile element class in the human genome being the SINE elements of size around 300 bp. Notably, the novel sequence insertion type carried a greater proportion of large insertions than other types, with 164 (19%) of the 869 novel sequences larger than 1,000 bp.

**Figure 3:**
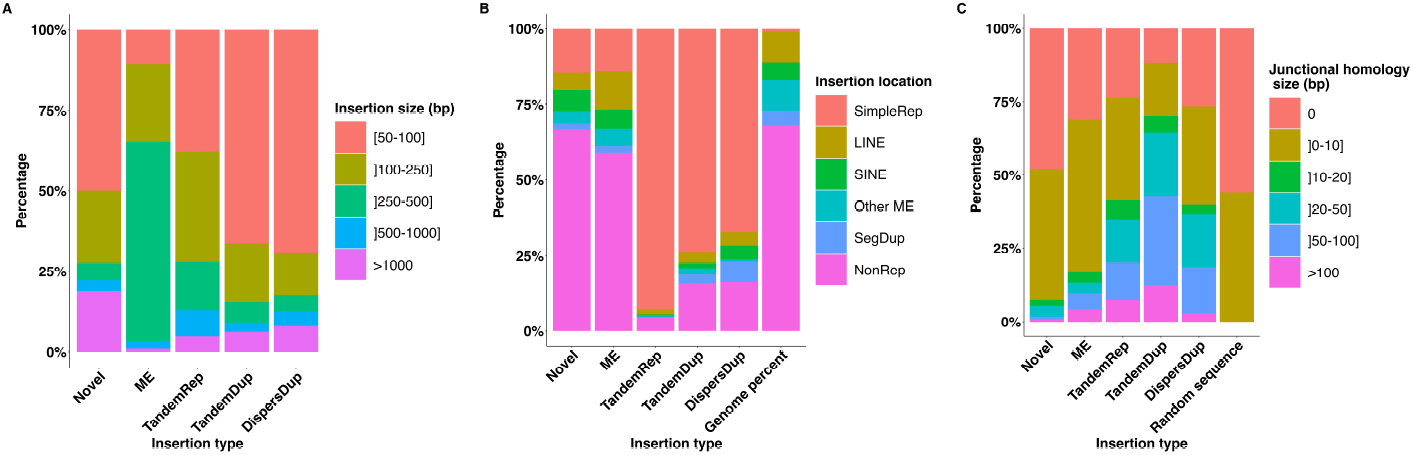
Proportions of insertion variant features according to the type of insertion. Proportions of classes of (A) insertion size, (B) insertion location, (C) homology size at the breakpoint according to the type of insertion in the NA19240 callset. Abbreviations: SimpleRep for simple repeat, ME for mobile element, TandemRep for tandem repeat, TandemDup for tandem duplication, DispersDup for dispersed duplication.

#### Characterization of insertion locations in the genome

We then characterized the insertions based on the genomic context of their insertion site. We investigated in particular genomic features that might make read mapping and SV calling difficult, such as the repetitive content. A strong over-representation was found in regions annotated as simple repeats, with 9,675 (70%) of the annotated insertions located in these regions that only represent 1.2% of the genome (Figure 2 C). The preferred genomic context of insertions varied between insertion types (Figure 3 B). 8,047 (92%) tandem repeats, 723 (73%) tandem duplications and 519 (63%) dispersed duplications were found in simple repeat regions. Conversely, 580 (67%) novel sequence insertions and 1,383 (56%) mobile element insertions were located in other regions. We did not find a higher rate of insertions within exonic, intronic or intergenic regions compared to a uniform distribution along the genome.

#### Junctional homology

We systematically compared the insertion site junction sequences with the inserted sequence extremities to identify stretches of identical or nearly identical sequences, here-after called junctional homologies as in [16] (see Material and Methods). Overall 5,119 (38%) insertions showed junctional homologies larger than 10 bp (Figure 2 D). This proportion is greater than the one obtained with random sequence insertions, the largest observed junctional homology being of 7 bp among 2,000 randomly simulated insertions (see Material and Methods). All insertion types carried junctional homologies greater than expected with random sequences. Tandem duplications and tandem repeats are the types with the greatest junctional homologies, with 428 (43%) tandem duplications and 1,751 (20%) tandem repeats that were identified with a junctional homology larger than 50 bp (Figure 3 C).

### Comparison with other individual callsets

These observations were performed on the NA19240 individual callset. Hence, we asked whether they could be recurrent across individuals from various genetic backgrounds. We first considered the two other individuals of the Chaisson et al study, namely HG00514 (14,363 insertions), son of a Han Chinese (CHS) trio, and HG00733 (15,476 insertions), son of a Puerto Rican (PUR) trio. These callsets were obtained with the same sequencing technologies and SV calling methodologies as for the NA19240 individual. Then, we analyzed a callset obtained by a different study, namely the SV callset for individual HG002 (13,541 insertions) provided by the Genome in a Bottle (GiaB) Consortium [19]. In this study, Zook and colleagues also used multiple sequencing technologies and SV calling methods to achieve a high confidence insertion and deletion callset (see Supplementary Table 1 for a summary of the technologies and methods used for all the callsets). Before comparing insertion features between callsets, we first checked whether they contained different variants. Using a rough estimation of shared variants, we identified only 1,169 insertion sites common to the four callsets within a 1 kb size window. On average 3,344 insertions were shared between two given callsets, and overall, more than 55% of the studied insertions were specific to a given callset. The distributions of insertion types, sizes, locations and junctional homology sizes were similar between the three individuals of the Chaisson et al study and the GiaB callset (Figure 2).

### Short-read-based recall

In order to investigate whether the previously described insertion features impacted the recall of short-read-based (SR-based) SV callers, we reproduced our previous analysis according to the technology involved in the variant call as annotated in the callsets. For the individual NA19240, 2,363 (17%) insertion variants were comforted by SR-based SV callers. As shown in Figure 4, this SR-based recall was highly heterogeneous with respect to the previously described insertion features. Each described feature in this work (ie. nature and size of the inserted sequence, insertion site genomic context and junctional homologies) impacted the SR-based recall. As shown in Figure 4 A, insertions larger than 500 bp were poorly discovered by SR-based methods (< 3 %). An increased SR-based recall for the 250-500 bp insertion size class corresponded to mobile element insertions. The greatest difference in SR-based recall was observed among the insertion types: 1,410 (57%) mobile elements and 342 (40%) novel sequence insertions could be detected with SR-based SV callers compared to only 87 (9%) tandem duplications, 484 (6%) tandem repeats and 40 (5%) dispersed duplications (Figure 4 B).

**Figure 4:**
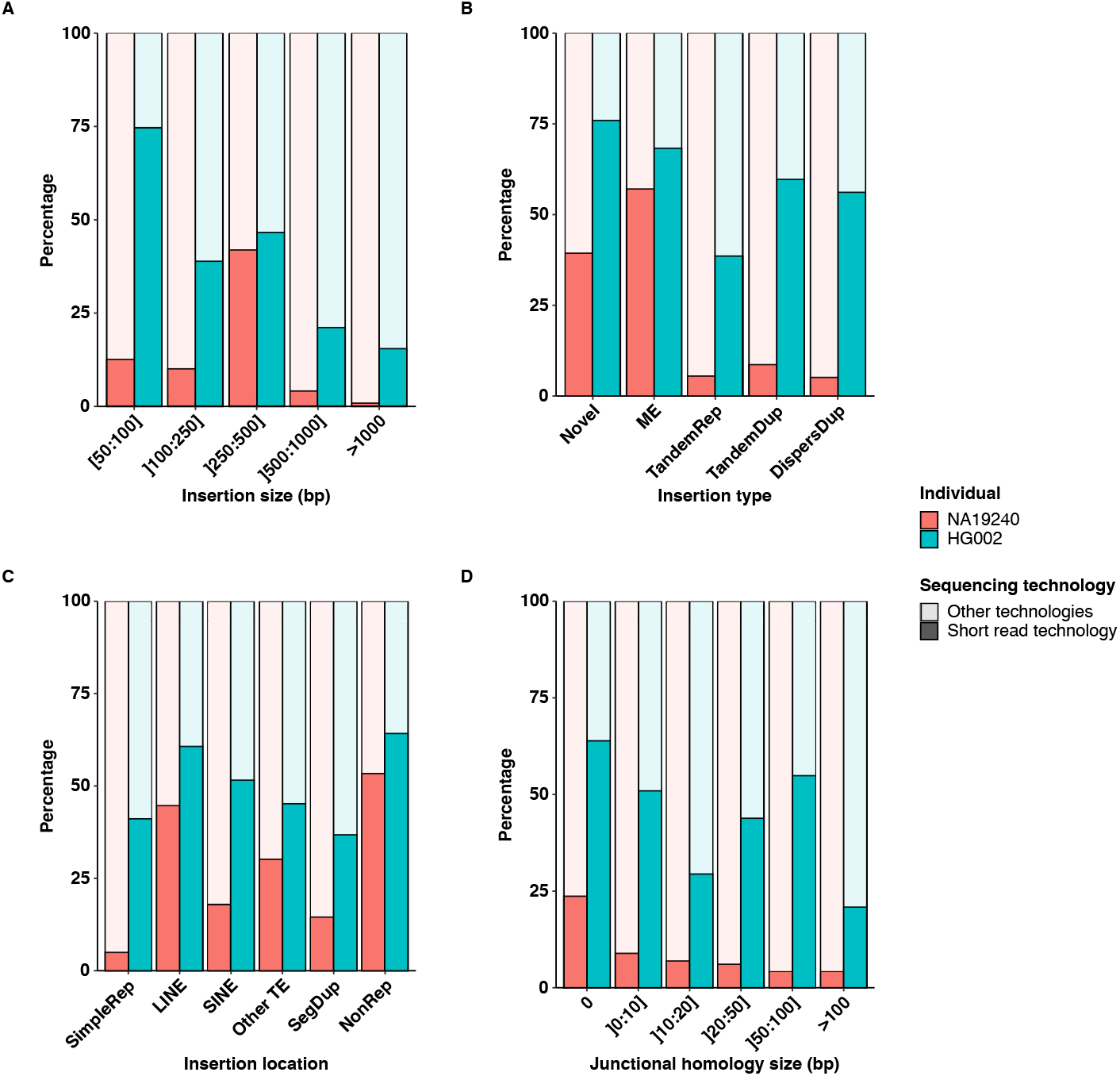
Proportions of SR-based insertion discoveries according to insertion features. Proportions of SR-based insertion discoveries (Short read technology) according to (A) insertion size, (B) insertion type, (C) insertion location and (D) homology size at the breakpoint, in the NA19240 and HG002 callsets.

The variations of the SR-based recall with respect to insertion features were very similar between the three studied individuals from the Chaisson et al. study (Supplementary Figure 2). However, the same comparison across two different studies with different methodologies was much more contrasted. Firstly, overall around twice as much insertions in proportion could be detected by SR-based methods in the GiaB study compared to the Chaisson et al study (SR-based recalls of 37 % and 17 % for HG002 and NA19240 callsets respectively). Secondly, the SR-based recall was much more homogeneous with respect to insertion features in the GiaB callset (Figure 4). The feature showing the most impact was the insertion size with a decrease of the SR-based recall with the insertion size, reaching below 2 % for insertions larger than 1 Kb (Figure 4 A). Similarly to the NA19240 callset, tandem repeats appeared more difficult to discover with SR-based methods, but to a lesser extent in the GiaB callset (Figure 4 B). Insertions located in simple repeats were less discovered using SR-based methods but this SR-based recall of 40 % remained higher than for NA19240 where it only reached 5 % on these locations (Figure 4 C). In contrast to the Chaisson et al study, junctional homology of the insertions of individual HG002 did not influence its SR-based recall (Figure 4 D).

### Using simulations to investigate the factors impacting the insertion calling recall

In real insertion callsets, most of the previously identified factors impacting SV discovery are intertwined. In order to quantify the impact of each factor independently, we produced various simulated datasets of 2×150 bp reads at 40x coverage, containing each 200 homozygous insertion variants on the human chromosome 3. As a baseline, we simulated 250 bp novel sequences taken from *Saccharomyces cerevisiae* exonic sequences inserted inside human exons. This is meant to represent the easiest type of insertions to detect. Then, we considered four scenarios of simulations, where only one of the four previously studied factors is changed at a time with respect to the baseline simulation.

Four insertion variant callers were evaluated on these datasets. They were chosen according to their good performances in recent benchmarks [7] and to maximise the methodological diversity. GRIDSS [11], Manta [20] and SvABA [6] are based on a first mapping step to the reference genome, contrary to MindTheGap (MTG) [10] which uses solely an assembly data structure (the De Bruijn graph). Two types of recall were computed depending on the precision and information given for each call: *insertion-site only* recall only evaluated if an insertion was called at an expected genomic position regardless of the predicted size or inserted sequence. As a more stringent evaluation, the *sequence-resolved* recall considered as true positives only those insertion calls having a correct genomic position and whose inserted sequence was very similar to the simulated one (>90 % sequence identity and +/- 10% insertion size).

#### Factors impacting insertion site recall

Recalls of insertion sites for all four methods are presented for the different simulated datasets in Table 1. On the baseline simulation, all tools succeeded to detect 100 % of simulated insertions, except for GRIDSS with 81 % of recall. The size of the inserted sequence impacted the recall of the insertion sites for most tools, except MindTheGap. GRIDSS was challenged by small insertions (50 bp) whereas Manta and SvABA had more issues with large insertions. The most extreme behavior was observed for SvABA which was not able to find the insertion sites of any of the simulated novel sequences larger than 500 bp.

**Table 1:** Insertion site recall of several short-read insertion callers according to different simulation scenarios. Cells of the table are colored according to the variation of the recall value of the given tool with respect to the recall obtained with the baseline simulation (first line, colored in blue): cells in red show a loss of recall >10%, cells in grey show no difference compared to baseline recall at +/- 10%. MTG: MindTheGap.

|  |  | Insertion site only recall (%) |  |  |  |
| --- | --- | --- | --- | --- | --- |
|  |  | GRIDSS | Manta | SvABA | MTG |
| **Baseline simulation:** 250 bp novel seq. in exons |  | 83 | 100 | 100 | 100 |
| **Scenario 1**<br>**Insertion**<br>**size** | 50 bp | 56 | 100 | 100 | 100 |
|  | 500 bp | 100 | 86 | 0 | 99 |
|  | 1,000 bp | 100 | 88 | 0 | 98 |
| **Scenario 2**<br>**Insertion**<br>**type** | Dispersed duplication | 100 | 1 | 100 | 96 |
|  | Tandem duplication | 100 | 100 | 100 | 0 |
|  | Mobile element | 100 | 2 | 100 | 58 |
|  | Tandem repeat (6 bp pattern) | 100 | 90 | 1 | 0 |
|  | Tandem repeat (25 bp pattern) | 99 | 66 | 0 | 2 |
| **Scenario 3**<br>**Junctional**<br>**homology** | 10 bp | 100 | 100 | 96 | 0 |
|  | 20 bp | 100 | 100 | 85 | 0 |
|  | 50 bp | 77 | 68 | 12 | 0 |
|  | 100 bp | 100 | 22 | 49 | 0 |
|  | 150 bp | 100 | 0 | 100 | 0 |
| **Scenario 4**<br>**Genomic**<br>**location** | Non repeat | 83 | 100 | 94 | 83 |
|  | Simple repeat (<300 bp) | 82 | 100 | 100 | 73 |
|  | Simple repeat (>300 bp) | 87 | 94 | 95 | 58 |
|  | SINE | 90 | 100 | 99 | 53 |
|  | LINE | 80 | 100 | 97 | 90 |
|  | Clustered insertions (<150 bp) | 85 | 85 | 2 | 77 |
|  | Real locations | 84 | 80 | 71 | 38 |
| **Scenario 5: real insertions at real locations** |  | 39 | 35 | 44 | 6 |

When simulating various insertion types, GRIDSS was the only tool whose recall was not negatively impacted. Manta could not find any type of dispersed duplications and showed a lower recall to detect tandem repeats with 25 bp size seeds. MindTheGap was unable to detect any type of tandem duplications and found only 58 % of mobile element insertions. SvABA was not able to detect any tandem repeat insertion but was able to detect all dispersed and tandem duplications and mobile elements.

Concerning junctional homology, the tools showed contrasting behaviors. GRIDSS was the only tool unaffected by the presence and size of repeated sequence at the insertion junctions. On the contrary, MindTheGap was the most impacted by junctional homology, being unable to detect insertions with homology at any tested size. This feature is actually controlled by a parameter of MindTheGap, increasing the max-repeat parameter value to 15 bp (default: 5bp), MindTheGap discovered 99 % of the insertion sites with 10 bp junctional homomolgies.

The recall of Manta decreased with the size of junctional homologies, whereas SvABA handled small (less than 20 bp) or very large (150 bp) junctional homologies but was affected by medium sizes. Concerning the impact of the genomic context of insertions, no loss of recall was observed in non repeated locations for GRIDSS and Manta. SvABA and MindTheGap displayed a 6 % and 17 % loss of recall respectively in these locations. Alignment-based SV callers showed no change in recall in small simple repeat (<300 bp), SINE and LINE locations. Manta and SvABA recalls lost 5 to 6 % of recall in simple repeat regions larger than the insert size (>300 bp). MindTheGap lost 42 and 47 % of recall in large simple repeat and SINE location simulations. Simulating insertions close to each other on the genome, at less than 150 bp, reduced the recall of SvABA (−98%), MindTheGap (−33 %) and Manta (−15%). Manta and MindTheGap obtained their smallest recall for the simulated dataset with 200 insertion sites sampled from the real locations of NA19240 chromosome 3 insertions.

Finally, when simulating the 889 insertions of NA19240 callset located on chromosome 3, with their reported inserted sequence at their real locations as described in the variant calling file (scenario 5), the recall of all tools dropped to less than 44 %, reaching for many tools their lowest values among the different simulated datasets. This was particularly marked for GRIDSS whose recall was greater than 77 % in all simulated scenarios, but achieved only 39 % on this simulation.

#### Impact of quality filtering

Previous results were computed using only the calls assessed with sufficient quality by each tool and annotated as PASS in the FILTER field of the VCF file. Removing this quality filtering allowed to increase the recall mainly for GRIDSS and SvABA (see Supplementary Table 3). Remarkably, GRIDSS reached a 100% recall on almost every scenario, except the scenario simulating the real insertions where still a 35% loss of recall was observed (Supplementary Table 3). These differences indicated that a substantial amount of true positive insertions were detected but reported as low quality calls.

#### Sequence-resolution of predicted insertions

We then investigated whether the SV callers were also able to recover the full inserted sequences in the different simulation scenarios (Table 2). On the baseline simulation with 250 bp novel sequence insertions, every tools reported for almost all detected insertion sites a resolved and correct inserted sequence. However, these high sequence-resolved recalls dropped dramatically when deviating from the baseline scenario. Although the discovery of insertion sites was not much impacted by the insertion size, all tools but MindTheGap were not able to recover any of the inserted sequences when it was larger than 500 bp (Table 2). On the contrary, MindTheGap assembled correctly nearly all simulated novel sequences, even those of 1 Kb.

**Table 2:**
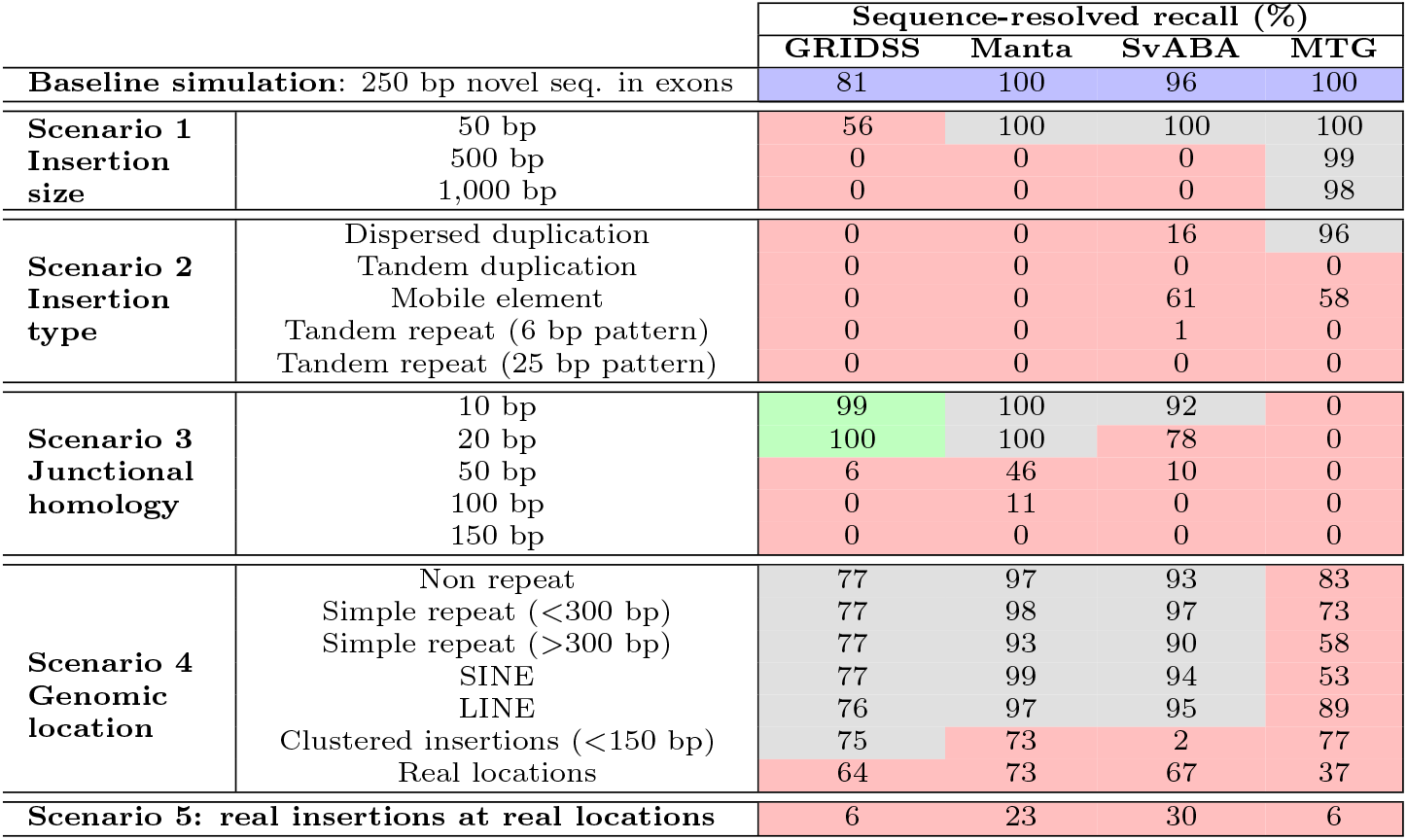
Sequence-resolved recall of several short-read insertion callers according to different simulation scenarios. Cells of the table are colored according to the variation of the recall value of the given tool with respect to the recall obtained with the baseline simulation (first line, colored in blue): cells in red show a loss of recall >10%, cells in grey show no difference compared to baseline recall at +/- 10%. MTG: MindTheGap.

Concerning the other insertion types, tools were not able to provide sequence resolved calls, except for MindTheGap and SvABA for some dispersed duplications and mobile element insertions (Table 2). In the case of tandem repeats, GRIDSS which detected all insertion sites, reported inserted sequences of at most 150 bp (instead of 250), corresponding to the simulated read size. The increase of junctional homology size reduced the sequence resolution of GRIDSS and SvABA. Insertions located in repeated regions were less resolved than in the baseline simulation for every tools. Finally, the sequence resolution of real insertions simulated at their real locations decreased compared to the insertion site recall, GRIDSS suffering the greatest loss (−33%).

#### False positive amount variations

The tools with the largest recalls were also the tools producing the largest amounts of false positive discoveries (in the order of several hundreds for GRIDSS and SvABA, see Supplementary Table 4). More surprisingly, the amount of false positives was not constant for most tools between the different simulation scenarios. It increased when simulated insertions presented a duplicative pattern (mobile element, dispersed duplication and junctional homologies above 50 bp). Removing the quality filter led to a large increase of the amount of false positive discoveries for GRIDSS and SvABA (5 to 17 times more respectively).

#### Unions and intersections of SV callers

A classical strategy to report SVs on real data is to reconcile several SV callsets keeping only variants that are similarly called by different SV callers. This strategy ensures a balance between true and false disovery rate. On the last simulation scenario, only 12 % of the insertion sites were validated by the three tools, GRIDSS, Manta and SvABA, and 39 % by at least two tools. However, the union of all three methods comprised 65 % of the real insertion sites, which represented an increase of 20 % of the best recall obtained by a single method (Figure 5).

**Figure 5:**
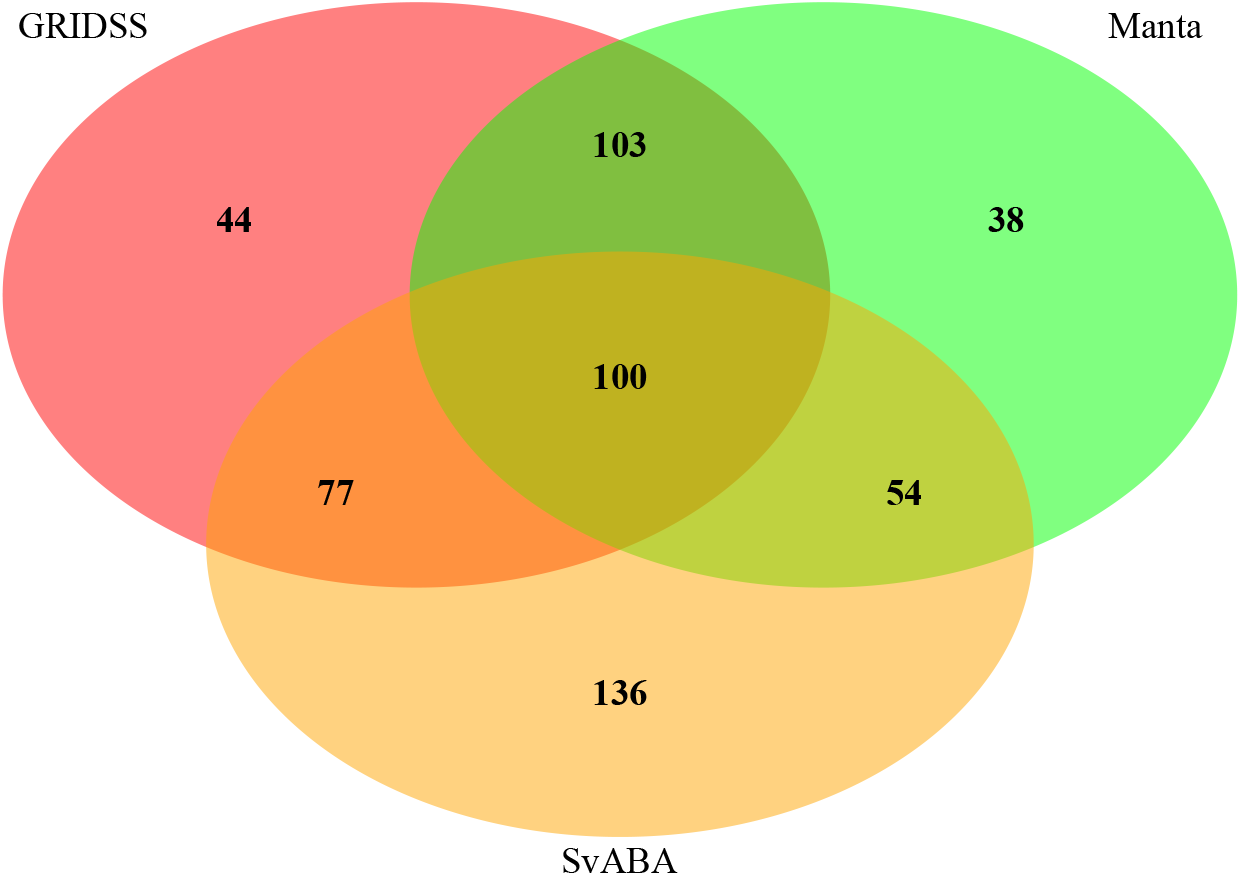
Intersections of true positive insertion callsets between different SV callers. Intersections of true positive insertion callsets between GRIDSS, SvABA and Manta on the scenario 5 simulation (real insertions at real locations). In this scenario, the 889 insertions located on the chromosome 3 from the NA19240 callset were simulated as described in the vcf file. Insertion calls were validated and compared based solely on the insertion site prediction.

## Discussion

The discovery of genomic variants is an important step towards the understanding of genetic diseases and species evolution [21, 22]. The detection of insertions too small (< 1kb) to be detected using comparative genomic hybridization array (CGH array) but larger than indel size (>50 bp) to be detected by the gold standard small variant discovery pipeline (GATK), remained a challenge with short read technology [4]. Thus these variations were poorly characterised in databases as compared to other SVs such as deletions. Numerous variant callers have been developed to overcome this issue but without resolving it [7]. Long read technologies or the crossing of various sequencing technologies overcome these limitations but are not affordable for many applications such as routine diagnosis of genetic diseases [18]. Thus, to improve current and future SR based SV callers, a better understanding of the actual insertion variants present in human populations is required.

We have presented here one of the most detailed and comprehensive analyses of actual insertion variants in the human genome looking for factors impacting their detection with short read re-sequencing data. This could be possible thanks to the publication of two exceptional SV callsets by Chaisson et al[18] and Zook et al (GiaB) [19]. These catalogs of insertions are considered as the most exhaustive for a given human individual and are qualified as gold standards thanks to their extensive validation by extensive and cross technology sequencing datasets. Unlike in the Chaisson et al study, the GiaB callset contained two categories of variants: 7,244 insertions that were reported with a higher confidence (*PASS* in the *FILTER* field) and 6,297 other insertions. As mentioned by the authors, the first category is likely to be biased towards easier to discover variants. Because we did not want to introduce this potential bias, and after checking that these two categories showed similar insertion feature distributions (see Supplementary Figure 1), we decided to conduct our analyses on the whole callset.

Not only, these catalogs of insertion variants are considered as the most exhaustive for a given human individual, but they are also the first sets with sequence-resolved events for any size and type of insertions. The fine resolution of the inserted sequences, present in these datasets, enabled us to propose a refined classification of insertion variants. In the two datasets, insertion types were not formally defined and the classifications differed between the datasets. Our classification allowed to normalize these heterogeneous annotations and was a direct application of variant definitions from the dbVar database which is based on the sequence ontology (SO). [12] We based our insertion type annotation on a minimal sequence coverage threshold, that was set to a relatively high value, 80%, in order to ensure a good specificity of our annotation. Increasing this value led to many more unassigned insertions, as the annotations were based on sequence alignments that were affected by potential remaining sequencing errors in the inserted sequences, polymorphism with the reference genome and the usage of alignment heuristics. If the amount of unassigned insertions decreased with the coverage threshold value, proportions of the different insertion types remained quite stable (Supplementary Table 2). Among the 12 % of unassigned insertions, some could correspond to a mixture of several insertion types, which particular case was not considered in this study. We did not define segmental duplications and CNVs as additional sub-types of dispersed duplications, as they are defined in the literature by their size (above 1 Kb), whose threshold has been arbitrarily set according to detection limitations of ancient technologies. As previously reported in the Chaisson et al and GiaB studies, we observed a highly heterogeneous distribution of insertion types and locations along the genome. The vast majority of insertions consisted in tandem repeats (63%) and most insertion sites were located in simple repeat regions (70%). These regions of low complexity, although representing a small proportion of the genome (1.2 %), are therefore a major source of inter-individual variability.

The sequence-resolution provided in these SV callsets also enabled us to analyze precisely the breakpoint junctions of each insertion variant. Junctional homology has been shown to be a frequent feature of SVs, that can be used to infer the rearrangement molecular mechanism [14, 15]. Although, it has been previously described for human SV callsets (around 2,000 SV breakpoints, including less than 400 insertions) [15], this is, to our knowledge, the first exhaustive quantification of junctional homology for such a large and almost complete set of insertions in a human individual. However, our measure of homology size is highly dependent on the callset precision of the insertion site location and of the inserted sequence. As SVs are often difficult to precisely localize, are subject to left-normalization processes, and their inserted sequences were mostly obtained from error-prone long reads, our measures may likely result in an under-estimation of the actual homology sizes. Despite these potential biases, our results show that real insertion variants harbour substantially larger junctional homologies than insertions that would be drawn randomly. Our measures allowed us to compare such feature between insertion types and all insertion types have been found to have a substantial proportion of variants with large junctional homologies (greater than 20 bp). Results showed also that large insertions tended to carry larger junctional homologies. As expected by their tandem nature, tandem repeats and tandem duplications had larger homology sizes than other insertion types. However, it was still smaller than their insertion size for many of them. The explanation for tandem repeat lies in their structure which is an amplification of a seed in the reference genome. Thus the largest homology size corresponded to the seed size presents at the right breakpoint (in case of left normalization). As for tandem duplications, the discordance between their annotation as tandem duplication and the smaller size of the detected junctional homology was related to the difference in the methods used to define the homology, where an exact match was required in the junctional homology case. This result suggests that many tandem duplications have small variations with their adjacent copy that break the exact match.

All the features of insertions characterized in our study (ie. nature and size of the inserted sequence, insertion site genomic context and junctional homologies) showed to impact the ability of SR-based SV callers to discover these variants, as defined by method annotations in the SV callsets. However, an important difference was observed between the two studies, with the GiaB study being able to detect with short reads twice as many insertions in proportion than in the Chaisson et al study. The difference in SR-based recalls between the two studies can certainly be explained by the different SR-based tool sets used and the different callset filtering and merging methodologies. The two studies used roughly the same number of SV-callers (13 and 15), but with a poor intersection: only one SV-caller (Manta) was common to both studies. Additionally, the method annotation of each variant is highly dependant on the study methodology to filter and merge the numerous callsets obtained for the same individual with different sequencing technologies and SV callers. For instance, it is not clear, if the presence of an SR-based tag for a given variant does necessarily mean that the latter can be discovered and sequence-resolved solely using short reads. However, both studies showed similar weaknesses to detect tandem repeats, large insertions and insertions located inside simple repeats. These observations are in-line with the already known difficulties of mapping short reads in such contexts.

These disparities between studies and the fact that most identified factors responsible of low SR-based recall are intertwined with one another in real insertion variants led us to pursue these investigations with simulated data. Our simulations did not aim at providing an exhaustive benchmark of SV callers but at identifying the precise genomic factors of insertion variants that prevent their correct discovery with short reads. As a consequence, we selected a small but diverse set of SV callers and we deliberately ran them with their default parameters. We based our selection of SV callers on a recent and comprehensive benchmark study by Kosugui et al [7]. SV callers selected in our study were chosen for their good performance in this benchmark, for their diversity of algorithms and for their ease of installation and usage. MindTheGap was not among the best insertion callers identified by Kosugui et al but was the only one not based on read mapping and using intensively de novo assembly with the whole read dataset.

Simulations remain a powerful approach to identify the strengths and weaknesses of SV callers but they were not meant to reflect perfectly real situations. In our simulations, several features may be far from the real complexity of human genome re-sequencing, such as some sequencing technology biases, the use of one chromosome instead of the whole genome, and the absence of other polymorphisms than insertion variants (SNPs, small indels and other SVs). As a consequence, the reported recalls are likely to be over-estimations of the ones obtained with real data. Although absolute values should be interpreted with caution, they can readily be compared between SV callers and between simulation scenarios. As a matter of fact, we often observed strong differences in recalls allowing to provide interesting insights in terms of impacting factors and SV caller behaviors. Our simulation protocol enabled to study each difficulty factor independently and highlighted the larger impact of insertion type compared to insertion location. However, all studied factors taken independently could not explain the whole loss of recall when simulating the real insertions at their real locations and there is probably an important synergetic effect of combining in a single insertion event several of the studied factors. For instance, the discovery of novel sequences in repeated regions was not a problem for almost every tested tools. However, the change of novel sequences to real inserted sequences, most of them corresponding to tandem repeats, reduced by half the recall of SV callers.

Our simulations revealed that junctional homologies as small as 10-20 bp impacted the recall of all tested tools. Such repeated sequences are likely to alter the mapping signature targeted by SV callers. Although such features of SV breakpoints and their relation to the molecular mechanisms generating SVs have long been described, they seem to be rarely taken into account in the design of SV caller algorithms. Our study of the real insertions showed that such junctional homology sizes are relatively common, with almost 40 % of insertions with junctional homologies larger than 10 bp. Therefore, SV callers algorithms would benefit from taking into account such properties of the breakpoints, that are likely to generate very specific signals in terms of read mapping.

One striking result of our simulations is the absence of sequence resolution for most of the simulated insertion features and most of the tested SV callers. In addition to the obvious loss of information about the variant event, this also limits the identification of the insertion type, the genotyping and the validation of the predicted call. As a matter of fact, we observed that most insertions regardless of their type and insertion genomic context were detectable but often not reported with a sufficient quality due to this lack of resolution. Furthermore, sequence resolution is essential for the comparison and genotyping of SVs in many individuals. As these tasks are the basis for association studies and medical diagnosis, efforts should be directed towards a better resolution of the sequence of these variants [8, 23]. Results obtained with the local assembly tool MindTheGap showed that the use of the whole read dataset allowed many insertions and even large ones to be assembled. The restriction to a small subset of reads to perform local assembly may therefore be the shortcoming of the other tested SV callers. Resolving the inserted sequence is possible to some extent, but tandem repeats larger than the read size will remain difficult to resolve with short reads technology.

In this study, we mainly focused on the recall, but we also reported the amount of false positive discoveries. We chose to compare absolute amounts of false positive discoveries, rather than using the precision or FDR metrics, as the latter are dependant of the amount of true positive discoveries. Interestingly, the amounts of false positives were affected by the type of the simulated insertions. The largest amounts were observed for duplicative insertions, for which some SV callers predicted variants not only at the insertion site but also at the locations of homologous copies of the inserted sequences.

Overall, the different SV callers did not performed well in every situation and in every aspects of insertion calling. Each caller showed its own strengths and weaknesses, often different from the other tools. Precisely identifying these in terms of insertion variant features and genomic contexts will enable each tool to be used to its best advantage. To do so, benchmark studies should take into account the wide variability of variant features that this present work has highlighted. Two recent SV benchmarks have raised awareness of the variability in the performances of SV callers depending on data sets and approaches [7, 9]. They looked at several factors that could be responsible for this variability. Technical factors (reads size, insert size and sequencing coverage) and biological factors (nearby SNVs or indels, genomic context, and variant size) showed to impact the recall of SV callers. However, the latter factors were analyzed for all SV types combined and none of these studies took into account the different types of insertion variants. Best practices for benchmarking small variant calling have been suggested based on gold standard callsets in high confidence regions, leaving structural variation in the fog [24]. However, it is precisely this type of variation that requires best practices for benchmarking and a standardization of annotation as they are harder to identify and report. We hope that the present fine characterization of gold standard human SV callsets will help in the development of better practices for benchmarking SV callers.

Advises to improve the detection using short read technology have already been described such as the careful combination of complementary SV callers [7]. Meta SV callers such as Meta-sv, Parliament2 or sv-callers reconcile SV calls produced by different SV callers [25, 26, 27]. However, only the calls that are discovered concordantly between different tools are returned. This strategy allows the precision to be increased, but at the expense of the recall. Our simulations showed that the intersection of only three SV callers reduced the recall of 30 %, whereas taking their union could increase the recall by at least 20 %. Considering unions of callsets would require a careful control of false positive rates. A better control could probably be achieved with sequence-resolved variants and by taking into account the observed characteristics of the different insertion types. Another alternative, less described, could be the use of dedicated tools for each type of insertion, instead of using only general-purpose SV callers. Among them, Expansion Hunter has been designed to detect tandem repeats, Pamir and Popins for novel insertions and TARDIS for large duplications [28, 29, 30, 31].

## Conclusion

In this work, we produced a detailed characterization of the insertion variants in a given human individual. We identified many factors of human insertion variants that explain their low recall with SR-based SV callers, including complex insertion types, difficult genomic contexts, large insertion sizes and junctional homologies at the breakpoints. The significant variability in the characteristics of the insertion variants, as well as the fact that all difficulties were handled differently by the different tested SV callers, call for a better characterization and comparison of SV callers according to the targeted variant features. This better understanding will certainly allow SV calling algorithms to be combined and used in their best conditions. Such improvements are eagerly awaited by society for the development of personalised medicine and the resolution of diagnostic bottlenecks for many rare diseases.

## Material and methods

### Data origin

SV callsets of the following individuals were downloaded from the following links:

**NA19240:** ftp://ftp.ncbi.nlm.nih.gov/pub/dbVar/data/Homo_sapiens/by_study/genotype/nstd152/NA19240.BIP-unified.vcf.gz.
**HG00514:** ftp://ftp.ncbi.nlm.nih.gov/pub/dbVar/data/Homo_sapiens/by_study/genotype/nstd152/HG00514.BIP-unified.vcf.gz.
**HG00733:** ftp://ftp.ncbi.nlm.nih.gov/pub/dbVar/data/Homo_sapiens/by_study/genotype/nstd152/HG00733.BIP-unified.vcf.gz.
**HG002:** ftp://ftp-trace.ncbi.nlm.nih.gov/giab/ftp/data/AshkenazimTrio/analysis/NIST_SVs_Integration_v0.6/HG002_SVs_Tier1_v0.6.vcf.gz.

Only insertions that were sequence resolved (ie. with an inserted sequence entirely defined) and called also in at least one of the parents were kept. No filtering related to quality or coverage was applied. The human reference genome version for this study was Hg38 (GRCh38). To compare the callsets on the same reference genome, the HG002 callset produced on hs37d5 build was converted into Hg38 build using Picard [32], the hs37d5 to hg19 and the hg19 to hg38 chain files from GATK public chain files.

### Comparison of the callsets

As a rough estimation of the amount of shared insertion variants between callsets, insertion locations were compared regardless of the insertion type or sequence. Insertion variants located less than 1,000 bp apart from one another were considered as the same variant.

### Insertion type annotation

TandemRepeatFinder (TRF) was used to annotate tandem repeats within each inserted sequence [33]. Recommended parameters were used, except for the maximum expected TR length (-l) which was set to 6 millions. In order to annotate mobile elements in inserted sequences, we used dfam, one of the annotation tools of RepeatMasker [34]. Each inserted sequence was scanned by dfam with the standard HMM profile database of human mobile elements provided by the tool. For the annotation of dispersed duplications and the occurrence count of their copies in the reference genome, each inserted sequence was locally aligned against the Hg38 genome using Blat with default parameters [35]. Only the alignments with at least 90 % of sequence identity were kept. For the annotation of tandem duplications, the two sequences on either side of the insertion site and of the same size as the insertion were aligned against the inserted sequence using Blat.

We used a minimal sequence coverage threshold, *Min_cov_*, to annotate the insertions. To be assigned to a given sub-type, the inserted sequence had to contain at least one contiguous segment annotated with the corresponding type and covering at least *Min_cov_ %* of the inserted sequence. Novel sequence insertions were a special case where the contiguity of the annotation was not required: more than *Min_cov_* % of the inserted sequence should not be covered by any alignment with the reference genome nor with the mobile element reference sequences, nor contain tandem repeats. When several types fulfilled the minimal coverage requirement, only one type was assigned according to the decision tree described in Figure 1.

### Junctional homology detection

Junctional homology, as referred to and defined in [16], is a DNA sequence that has two identical or nearly identical copies at the junctions of the two genomic segments involved in the rearrangement. In the case of an insertion, a junctional homology is a sequence segment at the left (resp. right) side of the insertion site which is nearly identical to the end (resp. beginning) of the inserted sequence. Small junctional homologies (<10 bp on each side) were searched in a strict manner by scanning simultaneously the 10 bp sequence at the left (resp. right) side of the insertion site and the 10 bp end (resp. beginning) of the inserted sequence, counting the number of identical nucleotides starting from the insertion site until a mismatch is encountered. For larger homologies, both the 100 % identity and strict adjacency to the insertion site constraints were relaxed. We used the local alignments between the breakpoint junctions and the inserted sequence that were previously obtained with BLAT. Only the alignments with at least 90 % identity and occurring at a maximum of 10 bp before (resp. after) the insertion site and at a maximum of 10 bp from the end (resp. beginning) of the inserted sequence were retained. In case of multiple candidates hits at one side of the junction, the one located at the closest position from the extremities was kept. If homologies (small or large) were found at both sides of the junction, the homology size was obtained by summing both homology sizes after removing potential overlap on the inserted sequence. To compute the expected distribution of junctional homology sizes that could be observed by chance, we generated 2,000 random insertions on the human chromosome 3 sequence. Inserted sequences were generated by concatenating 250 nucleotides sampled uniformly on the A,C,G,T alphabet. The insertion sites were sampled uniformly along the chromosome 3 sequence. Junctional homology sizes of these random insertions were identified using the same previously described methodology as for real insertions.

### Genomic context characterization

To study the genomic context of insertions, we used the repeat content annotations of RepeatMasker from the UCSC genome browser for the Hg38 genome and the gene annotations from the Gencode v32 [36, 37, 38, 39].

### SR-based recall of the gold standard callsets

Each callset was partitioned in two parts based on the discovery technology. The first part, referred as *Short read technology*, contained insertion calls that carried the Illumina (short reads) tag or a SR-based caller tag. For Chaisson et al callsets (NA19240, HG00514 and HG0733), the selection was performed on the vcf *INFO* section and the *UNION* variable. The *UNION* variable can take three potential values, *Pacbio,Bionano* or *Illumina*, that corresponded to the sequencing technology allowing the variant to be discovered. For the GiaB callset (HG002), insertions that could be discovered with short reads were identified by the *Ill* tag contained in the *ID* or *INFO* sections of the vcf file. Insertion calls that were labelled *Ill* only with refining methodologies and not any discovery methodologies were not taken into account for the *Short read technology* part. The second part, referred as *Other technologies*, contained all the remaining insertions. It should be noted that all insertion calls in the first part carried also at least one long read technology tag and were not discovered using only short read technology.

### Simulations

Twenty two sequencing datasets were simulated to characterize the impact of the different insertion features on SR-based insertion variant calling. Each dataset was obtained by altering the human chromosome 3 with 200 insertions. Sequencing reads were generated using ART with the following parameters: 2×150 bp reads, at 40 X coverage, with insert size of 300 bp on average and 20 bp standard deviation [40].

#### Baseline simulation

The simulation referred as the baseline was meant to represent the easiest type of insertions to detect, where inserted sequences contained very few repeats and are novel in the genome, the genomic context of insertion was also simple and repeat-free, and breakpoint junctions did not have any homology. To do so, we simulated 250 bp novel sequence insertions located in exons without any homology at the breakpoint junctions. Novel sequences were extracted from randomly chosen exonic regions of the *Saccharomyces cerevisae* genome.

#### Scenario 1: varying the insertion size

Insertion locations used in the baseline simulation were kept and the 200 inserted sequences were alternatively replaced by sequences extracted from *Saccharomyces cerevisae* exons of 3 different sizes: 50, 500, and 1,000 bp.

#### Scenario 2: varying the insertion type

Insertion locations were identical to the baseline simulation, but the 250 bp inserted sequences were alternatively replaced by dispersed duplications, tandem repeats, tandem duplications and mobile elements. Two types of tandem repeats were simulated, with a pattern size of 6 bp or 25 bp, the pattern originating from the left breakpoint junction. As mobile elements, 200 Alu mobile element sequences with a size ranging between 200 and 300 bp were randomly extracted from the human genome based on the RepeatMasker annotation. Tandem duplications were generated by duplicating the 250 bp right breakpoint sequence. The inserted sequences of simulated dispersed duplications were extracted from exons of the chromosome 3.

#### Scenario 3: varying the junctional homology size

The 250 bp insertion sequences produced in the baseline simulation were altered with junctional homology. To simulate junctional homologies, we replaced the *X* first bases of each insertion with the same size sequence originating from the right breakpoint sequence. We simulated five junctional homology sizes (*X* value): 10, 20, 50, 100 and 150 bp.

#### Scenario 4: varying the genomic context of insertion

The 250 bp insertions from the baseline simulation were alternatively inserted in specific genomic contexts: either inside different types of mobile elements, namely SINEs and LINEs, in small (<300 bp) and large (>300 bp) simple repeats or in other regions not annotated by RepeatMasker (non repeated). A dataset with closely located variants was simulated by adding insertions closed to the insertions simulated in the baseline scenario. The distance between insertions varied uniformly from 5 to 150 bp. Finally, the simulation referred as “real locations” was simulated by relocating the 200 insertions from the baseline scenario to locations uniformly sampled from the 889 NA19240 insertions reported in the chromosome 3.

#### Scenario 5: Real insertions at real locations

The 889 insertions located on the chromosome 3 from the NA19240 callset were simulated as described in the vcf file.

### Insertion calling and benchmarking

Simulated reads were aligned with bwa against the hg38 reference genome, and read duplicates were marked with samblaster v.0.1.24 and converted into bam file with samtools v1.6 [41, 42]. Bam index and reference dictionary were obtained by picard tools v2.18.2 [32]. GRIDSS v2.8.0, Manta v1.6.0, MindTheGap v2.2.1 and SvABA v1.1.0 were all run using recommended, or otherwise default, parameters [11, 20, 10, 6]. Only “PASS” insertions, that were larger than 50 bp, were kept for the recall and false positive calculation. Two types of recalls were computed depending on the precision and information given for each call: insertion-site only recall and sequence-resolved recall. The insertion-site only recall was assessed solely based on the insertion site location prediction with a 10 bp margin around the expected location. As a more stringent evaluation, the sequence-resolved recall took also into account the inserted sequence. When it was reported, the inserted sequence had to share at least 90 % of sequence identity to the simulated one and had to have a similar size of +/- 10%, to be considered as a true positive. In case of absence of alternative sequence in the vcf file but the provided annotation of the event allowed us to extract the insertion sequence from the reference genome (for instance for dispersed duplication with the duplicated copy coordinates), it was evaluated similarly as for alternative sequences. Recall was computed as the ratio between the amount of true positive discoveries and the amount of simulated insertions.

### Code availability

Custom scripts used for the characterization of insertion callsets are freely available at https://github.com/WesDe/DeepAn. Scripts developed to simulate particular insertion variants and for the validation of vcf files are freely available at https://github.com/WesDe/InserSim.

## Competing interests

The authors declare that they have no competing interests.

## Author’s contributions

WD, JT and CL conceived the study. WD developed the analysis and simulation scripts and carried out the analysis of the results. All authors contributed to the writing and read and approved the final manuscript.

## Acknowledgements

We are thankful to the Genouest bioinformatics platform, computations have been made possible thanks to the resources of the Genouest infrastructure.

## Notes

### Competing Interest Statement

The authors have declared no competing interest.

