## Supplementary Material for "Towards a better understanding of the low recall of insertion variants with short-read based variant callers"

June 2020

[1] Univ Rennes, CNRS, Inria, IRISA - UMR 6074, F-35000, Rennes, France

[2] Unité de Génétique Clinique,Pôle Couple Enfant, CHU de Grenoble Site Nord-Hôpital Couple-Enfant, 38043, Grenoble, France

### 1 Characteristics of the studied callsets

| Study | Individual | Sequencing technology | Coverage | SV callers |
|---|---|---|---|---|
| Chaisson et al 2019 [?] | NA19240<br>HG00514<br>HG00733 | Illumina short insert | 77 | dCGH, Delly, GenomeStrip, NovoBreak,Pindel, retroCNV, SVelter, VH, Wham, Lumpy, ForestSV, Manta, MELT, Tardis_MEI, liWGS |
| | | Illumina liWGS | 179 | |
| | | Illumina 7kbp JMP | 37 | |
| | | 10X Chromium | 245 | No dedicated SV caller Home made strategy based on haplotype assembly and alignment on reference genome |
| | | BioNanoGenomics | 113 | |
| | | Tru-Seq SLR | 4 | |
| | | Strand-Seq | 7 | |
| | | Hi-C | 17 | |
| | | PacBio | 38 | |
| | | Oxford Nanopore (HG00733) | 19 | |
| Zook et al 2019 [?] | HG002 | Illumina HiSeq | 300 | Spirale Genetics tools, GATK-HC,Freebayes, Fermikits, MetaSV, TNscope, Scalpel, SvABA, Krunch, Cortex,Manta, Seven Graph Bridge Refinement |
| | | 10X Genomics | 86 | LongRanger |
| | | Complete Genomics | 100 | CGATools |
| | | PacBio | 44 | PbSv |
| | | | | Hybrid : HySA, BreakScan |

**Table 1: Sequencing technologies, sequencing coverage and SV callers used to generate the four high confidence SV callsets that were studied in this work.**

### 2 Characterization of insertion callsets

#### 2.1 Annotation of insertions

**Table 2: Annotation of the insertion callset of individual NA19240 according to the minimal sequence coverage threshold.** Bracketed values correspond to the category percentage among the annotated insertions.

| % Coverage | 100 | 95 | 80 | 60 | 40 |
|---|---|---|---|---|---|
| **New sequence** | 677 (10%) | 686 (6%) | 869 (6%) | 1,223 (8%) | 1,639 (11%) |
| **Mobile element** | 605 (9%) | 2,047 (17%) | 2,473 (18%) | 2,828 (19%) | 3,321 (22%) |
| **Tandem repeat** | 4,399 (65%) | 7,552 (62%) | 8,735 (63%) | 9,102 (62%) | 9,235 (61%) |
| **Tandem duplication** | 444 (7%) | 953 (8%) | 1,000 (7%) | 1,081 (7%) | 1,082 (7%) |
| **Dispersed duplication** | 486 (7%) | 816 (7%) | 774 (6%) | 767 (5%) | 713 (5%) |
| **Unassigned** | 8,890 | 3,456 | 1,843 | 1,046 | 473 |
| **% annotated** | 43.4 | 78.0 | 88.3 | 93.3 | 97.0 |

#### 2.2 Distributions of insertion variant features across several callsets

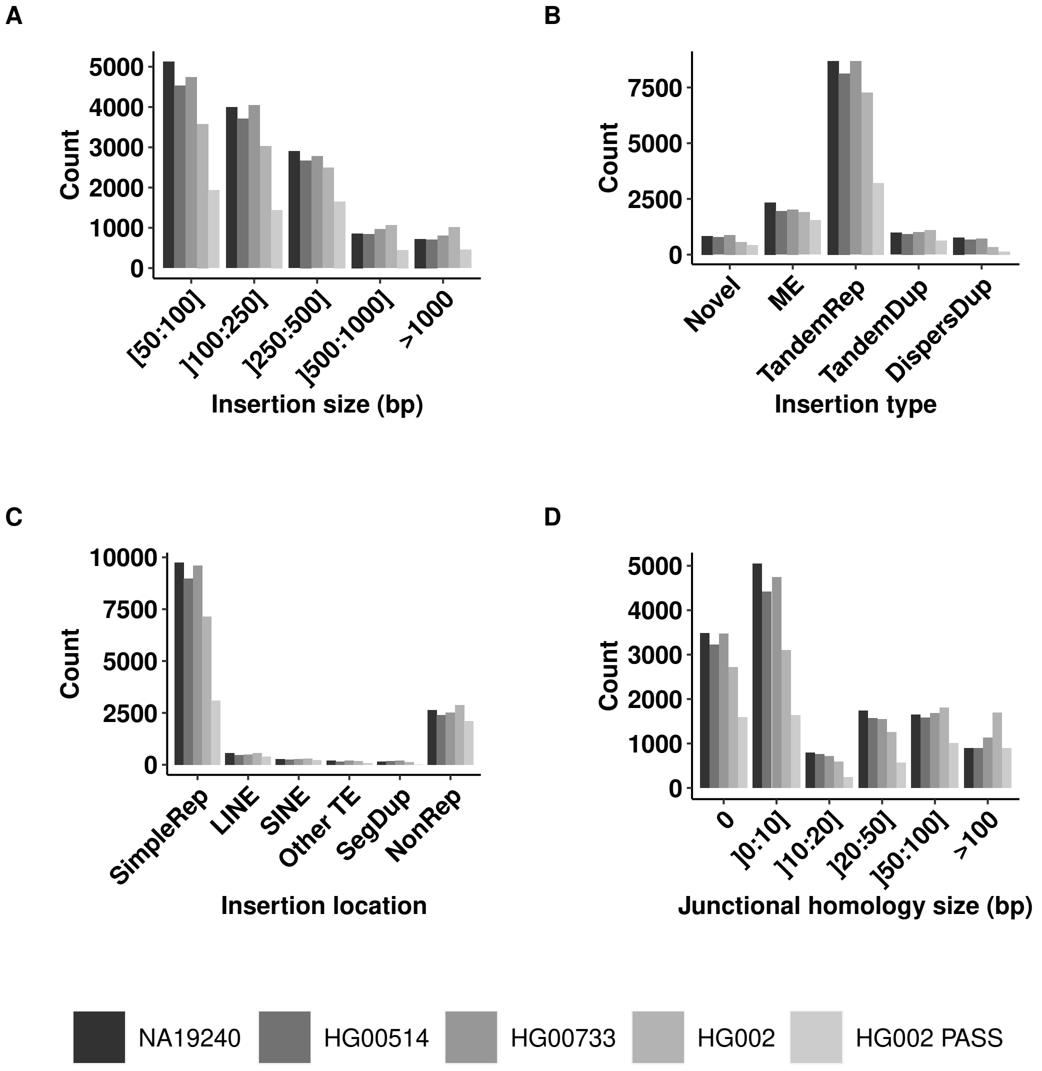

**Figure 1: Distributions of insertion variant features across several call sets.** Distributions of (A) insertion size, (B) insertion type, (C) repeated context of insertion and (D) homology size at the breakpoint for five insertion variant callsets. NA19240, HG00514, HG00733 refer to the three individual callsets of the Chaisson et al study. HG002 and HG002 PASS refer to the callset of the GiaB study, taking into account, respectivelly, the full callset or only the variants with PASS in the Filter field. Abbreviations: SimpleRep for simple repeat, ME for mobile element, TandemRep for tandem repeat, TandemDup for tandem duplication, DispersDup for dispersed duplication.

#### 2.3 Proportions of SR-based insertion discoveries according to insertion features

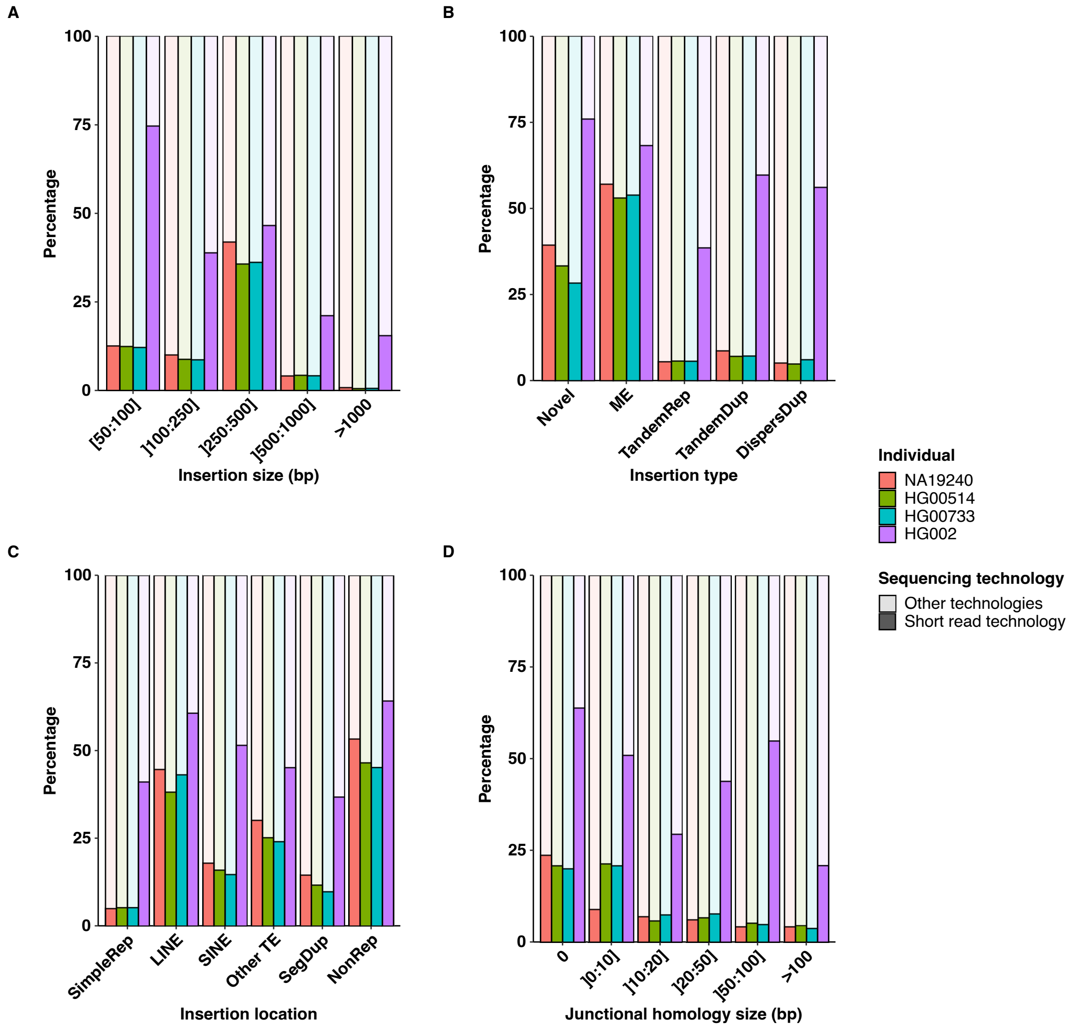

**Figure 2: Proportions of SR-based insertion discoveries according to insertion features for the four insertion callsets.** Proportions of SR-based insertion discoveries (Short read technology) according to (A) insertion size, (B) insertion type, (C) insertion location and (D) homology size at the breakpoint, in the three individual callsets from the Chaisson et al study (NA19240, HG00514, HG00733) and the HG002 callset from the GiaB study.

### 3 Simulation additional results

#### 3.1 Recall of SV callers without any quality filter

| | | Insertion site only recall - no quality filter (%) | | | |
|---|---|---|---|---|---|
| | | GRIDSS | Manta | SvABA | MindTheGap |
| **Baseline simulation**: 250 bp novel sequences in exons | | 100 | 100 | 100 | 100 |
| **Scenario 1 Insertion size** | 50 bp | 100 | 100 | 100 | 100 |
| | 500 bp | 100 | 86 | 6 | 99 |
| | 1,000 bp | 100 | 88 | 1 | 98 |
| **Scenario 2 Insertion type** | Dispersed duplication | 100 | 49 | 100 | 96 |
| | Tandem duplication | 100 | 100 | 100 | 0 |
| | Mobile element | 100 | 50 | 100 | 58 |
| | Tandem repeat (6 bp pattern) | 100 | 92 | 22 | 0 |
| | Tandem repeat (25 bp pattern) | 100 | 66 | 100 | 2 |
| **Scenario 3 Junctional homology** | 10 bp | 100 | 100 | 98 | 0 |
| | 20 bp | 100 | 100 | 89 | 0 |
| | 50 bp | 100 | 51 | 65 | 0 |
| | 100 bp | 100 | 12 | 100 | 0 |
| | 150 bp | 100 | 0 | 100 | 0 |
| **Scenario 4 Genomic location** | Non repeat | 100 | 100 | 98 | 83 |
| | Simple repeat (<300 bp) | 100 | 100 | 100 | 73 |
| | Simple repeat (>300 bp) | 99 | 94 | 100 | 58 |
| | SINE | 100 | 100 | 100 | 53 |
| | LINE | 100 | 100 | 100 | 90 |
| | Distance between insertion <150 bp | 100 | 85 | 77 | 77 |
| | Real locations | 96 | 81 | 90 | 38 |
| **Scenario 5: real insertions at real locations** | | 65 | 37 | 70 | 6 |

**Table 3: Insertion site recall of several SV callers without any quality filter applied.** For each SV caller, all predicted calls output in the final vcf file were taken into account regardless of their value in the FILTER field. Only the insertion site location is taken into account to compute the recall. Each line corresponds to a distinct simulation scenario. Cells of the table are colored according to the variation of the recall value of the given tool with respect to the recall obtained with the baseline simulation (first line, colored in blue): cells in red show a loss of recall >10%, cells in grey show no difference compared to baseline recall at +/- 10%.

#### 3.2 False positive amounts

**Table 4: Amounts of false positives called by the tested SV callers according to different simulation scenarios.** For each scenario involving several simulated datasets, the values indicate the minimal and maximal number of false positive predictions obtained over these datasets. Cells of the table are colored according to the variation of the FP amount of the given tool with respect to the amount obtained with the baseline simulation (first line, colored in blue): cells in red show a substantial increase of FP amount, cells in grey show small difference or a decrease of FP amount compared to the baseline simulation.

| | Amount of False positive calls | | | | | | | |
|---|---|---|---|---|---|---|---|---|
| | GRIDSS | | Manta | | SvABA | | MindTheGap | |
| | PASS | All | PASS | All | PASS | All | PASS | All |
| **Baseline simulation** | 0 | 151 | 2 | 2 | 6 | 84 | 19 | 19 |
| **Scenario 1: Insertion size** | 0 | 131 - 138 | 0 - 3 | 0 - 3 | 0 - 6 | 82 - 96 | 16 - 19 | 16 - 19 |
| **Scenario 2: Insertion type** | 3 - 400 | 233 - 591 | 0 - 18 | 0 - 201 | 4 - 451 | 92 - 1,157 | 17 - 19 | 17 - 19 |
| **Scenario 3: Junctional homology** | 2 - 9 | 128 - 163 | 0 - 4 | 0- 4 | 5 - 202 | 70 - 342 | 2 - 18 | 2 - 18 |
| **Scenario 4: Genomic location** | 0 - 4 | 143 - 166 | 0 - 5 | 0 - 5 | 4 - 13 | 74 - 643 | 16 - 19 | 16 - 19 |
| **Scenario 5: Real insertions** | 382 | 2,052 | 101 | 148 | 523 | 9,314 | 19 | 19 |
